# Reconstructing oral cavity tumor evolution through brush biopsy

**DOI:** 10.1101/2023.11.29.569216

**Authors:** Evit R. John, Tom Lesluyes, Toby M. Baker, Maxime Tarabichi, Ann Gillenwater, Jennifer R. Wang, Peter Van Loo, Xiao Zhao

## Abstract

Oral premalignant lesions (OPLs) with genomic alterations have a heightened risk of evolving into oral squamous cell carcinoma (OSCC). Currently, genomic data are obtained through invasive tissue biopsy. Brush biopsy has been utilized for diagnosing dysplasia but its effectiveness in reflecting the complete genomic landscape of OPLs remains uncertain. This study investigates the potential of brush biopsy samples in accurately reconstructing the genomic profile of OPLs. We analyzed single nucleotide variants (SNVs), copy number aberrations (CNAs), and subclonal structures in paired tissue and brush biopsy samples from a patient with both OPL and OSCC lesions. The results showed that brush biopsy can effectively reflect about 90% of SNVs and similar CNA profiles as those found in tissue biopsies. It was specific, as normal oral epithelium didn’t share these genomic alterations. Interestingly, brush biopsy revealed shared SNVs and CNAs between the distinct OPL and OSCC lesions, indicating a common ancestral origin. Subclonal reconstruction confirmed this shared ancestry, followed by divergent evolution of the lesions. These findings highlight the potential of brush biopsies in accurately representing the genomic profile of OPL and OSCC, proving useful in understanding tumor evolution.

## Introduction

Over 70 million people worldwide have oral premalignant lesions (OPLs) and these patients have a 2-5% risk of transformation to oral squamous cell carcinoma (OSCC) over a 5-year period^1^. Once transformed, overall survival is reduced to 60% and is associated with significant treatment morbidity^2^. The gold standard to assess transformation risk is based on histologic pathology. This process requires a biopsy of the suspect tissue, which is invasive and increases morbidity. In addition, there is variation between pathologist evaluators with unclear guidelines defining risk classification^3^.

While it is known that genomic alterations drive the progression of OPL to OSCC, there are no clinical biomarkers used to assess the risk of progression to invasive cancer. Losses of large chromosome fragments (e.g. 3p, 9p, 17p)^4^ or the presence of known drivers such as *TP53*^*5*^ have provided evidence associated with the risk of progression.

Brush biopsy has been used in cytology to identify dysplastic cells in oral lesions to diagnose OPLs^6,7^. While this serves as evidence that brush biopsies are capable of capturing tumor cells, it is unclear whether these tumor cells can provide an accurate representation of the genomic landscape of their parent tissue. If possible, brush biopsy may serve as a non-invasive strategy to study the genomic landscape of OPLs. This could lead to exciting opportunities to assess pathological mutations, clonal and sub-clonal structure, and reconstruct the evolutionary history of OPLs. Clinically, understanding the genomic landscape of OPLs through brush biopsy may provide prognostic and diagnostic utility.

In this proof-of-concept study, we compare whether brush biopsies of OPL and OSCC samples can recapitulate the genome-wide alterations found in their parent tumor tissue. Through this method, we provide the first evidence of an oral tumor seeding event to a distant site through a common ancestor and subsequent divergent evolution.

## Results and Discussion

Early detection and treatment of OSCC results in high overall survival rates and can often be managed with minimal surgical intervention^8^. While histopathology is the standard for assessing OPL risk, genomic alterations have shown potential for patient risk stratification, suggesting a potential role in prognosis. This proof-of-concept study evaluates whether brush biopsy can accurately reflect the genomic landscape of head and neck cancer, compared to traditional tissue biopsy. The patient studied had OSCC on the left tongue and OPL (moderate dysplasia) on the right, with these distinct lesions separated by normal epithelium (**Figure 1**). Tissue samples and brush biopsies were taken of the OPL, OSCC, and normal distant oral mucosa (buccal tissue). Whole-exome sequencing (WES) was performed to a median coverage of 145X.

**Figure 1:**
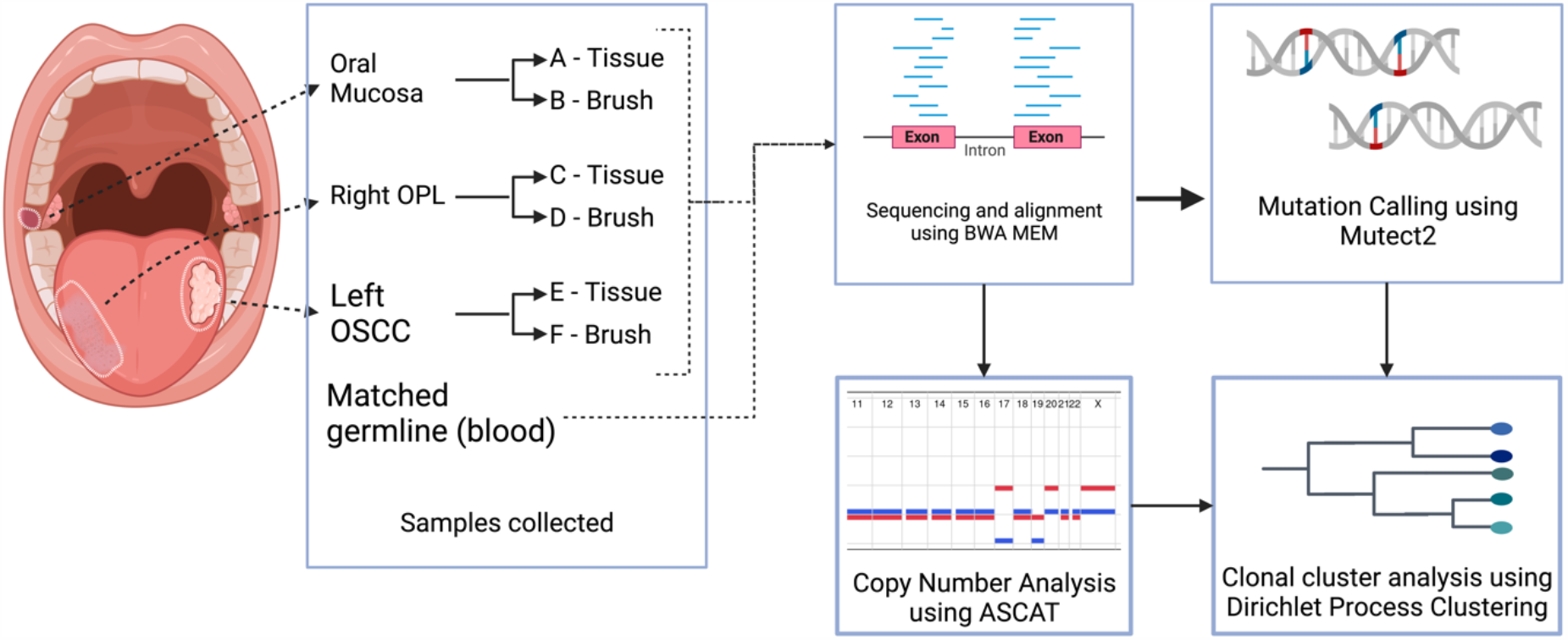
Workflow for subclonal reconstruction. Illustration of an oral premalignant lesion (OPL) and oral squamous cell carcinoma (OSCC) lesions and normal oral mucosa sampled from the same patient with both brush and tissue biopsies. The samples were aligned, and somatic mutations were called from the aligned reads. CNA profiles were constructed using ASCAT and SNV CCFs were then clustered to identify subclonal lineages in the samples and phylogenetic reconstruction was done.

Brush biopsies captured the majority of single nucleotide variants (SNVs) found in paired target tissue (**Figure 2**). The paired brush sample captured 85 out of 106 (80.2%) SNVs found in the normal oral tissue, 156 out of 185 (89%) in OPL, and 174 out of 189 (92%) SNVs found in the OSCC tissue. For oral mucosa and OSCC lesions, brush biopsy detected more mutations than tissue biopsy, with 167 (157%) more SNVs in the oral lesion and 17 (9.0%) more SNVs in OSCC. The substantial increase in mutation detection in oral mucosa brush samples might be attributed to sampling a broader, non-targeted area compared to the more focused tissue biopsy. However, in the case of the OPL lesion, tissue biopsy identified 10 (5.7%) mutations more than brush biopsy.

**Figure 2:**
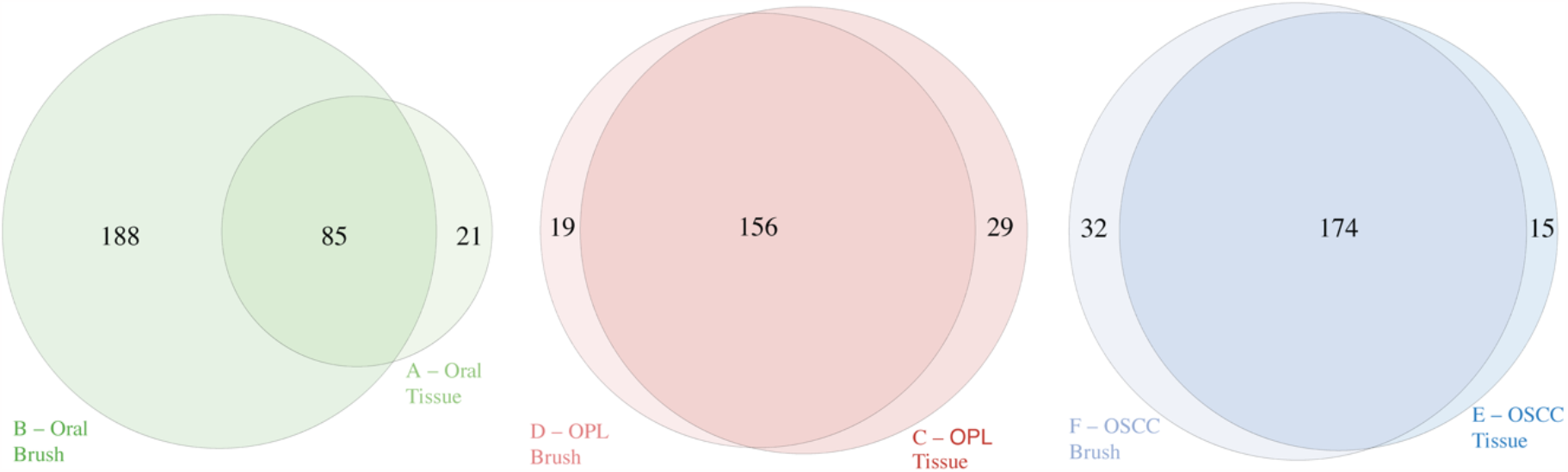
Mutations shared between the respective brush and tissue samples of each lesion.

To identify copy number alterations (CNA), we used Allele-Specific Copy Number Analysis of Tumors (ASCAT)^9^. Comparisons of CNA profiles between paired brush and tissue samples from each lesion revealed that brush biopsies accurately reflected the CNA of the corresponding tissue (**Figure 3**). In oral mucosa, OPL, and OSCC lesions, brush samples matched 98.7%, 81.6%, and 89.5% of the tissue samples’ allele-specific copy number calls, respectively. Notably, CNAs identified in normal epithelium tissue by brush biopsy were confirmed by tissue biopsy (**Figure 3**). Most significantly, both tissue and brush biopsies showed shared CNAs between the OPL and OSCC lesions, such as LOH of the short arm of chromosome 17^10,11^. Given the shared CNAs between the OPL and OSCC sample, the minimum event distance^12^ between ASCAT profiles was calculated and a copy number-based phylogenetic tree was reconstructed using MEDICC2^12^ (**Supplementary Figure 1**). This analysis showed a clear separation between normal epithelium and OPL/OSCC samples, with consistent clustering of the OPL and OSCC samples based on copy number events.

**Figure 3:**
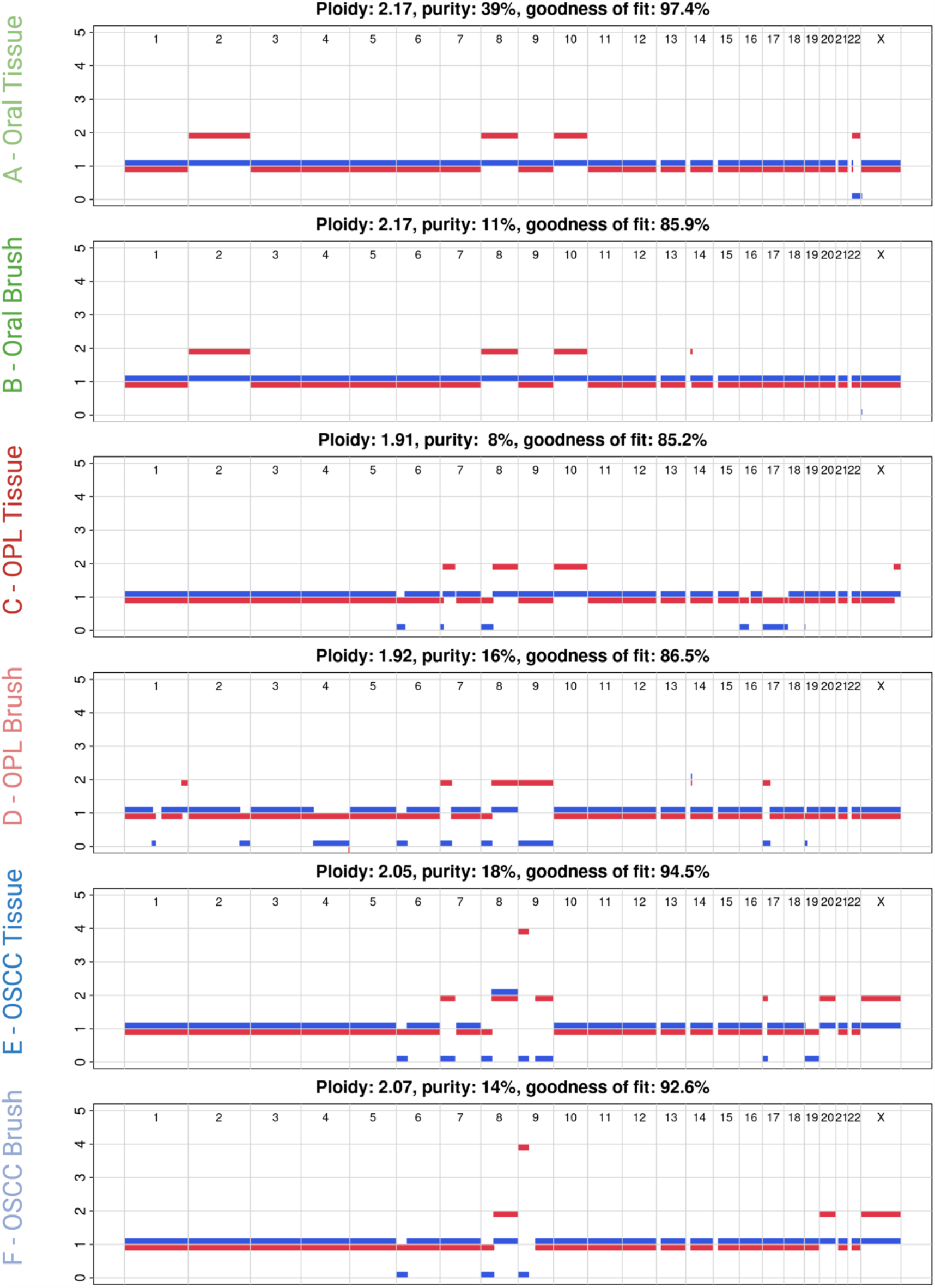
ASCAT profiles showing copy number alterations in each sample. Comparison of the tissue and brush samples show similar patterns in losses and gains in each lesion.

To identify clones and subclones, we used DPClust to cluster the SNVs based on their Cancer Cell Fraction (CCF) ^13^. The CCF is the proportion of tumor cells that carry each SNV and is calculated using SNV variant allele frequencies (VAF) corrected for purity, ploidy, and copy number. Shared and/or unique subclones were identified (**Figures 4 and 5a**). A consistent finding across all lesions was that brush biopsies captured all clonal and subclonal populations present in their corresponding tissue samples. Intriguingly, a unique subclonal population (cluster 7) was detected in the oral brush sample that was absent in the paired tissue, likely due to the broader sampling field of the brush biopsy. This is consistent with the larger number of SNVs detected in the brush sample (**Figure 2**). Paired samples from both OPL and OSCC showed a shared subclonal cluster, cluster 1, noting the presence of a common ancestor. This shared common ancestor was likely a seeding event rather than field cancerization, given that the lesions were found on the contralateral sides of the tongue with a large gap of normal epithelium between them. Following the shared cluster 1 ancestor, the OPL and OSCC samples separate with unique subclones, demonstrating evolutionary divergence. Finally, to evaluate the specificity of the brush biopsies, comparisons were made between brush samples from the OPL and OSCC lesions to the normal brush sample. The absence of shared SNVs or clonal/subclonal populations with the normal brush sample showed that the brush biopsies specifically captured the genomes of their target tissues (**Figure 4 and 5a**).

**Figure 4:**
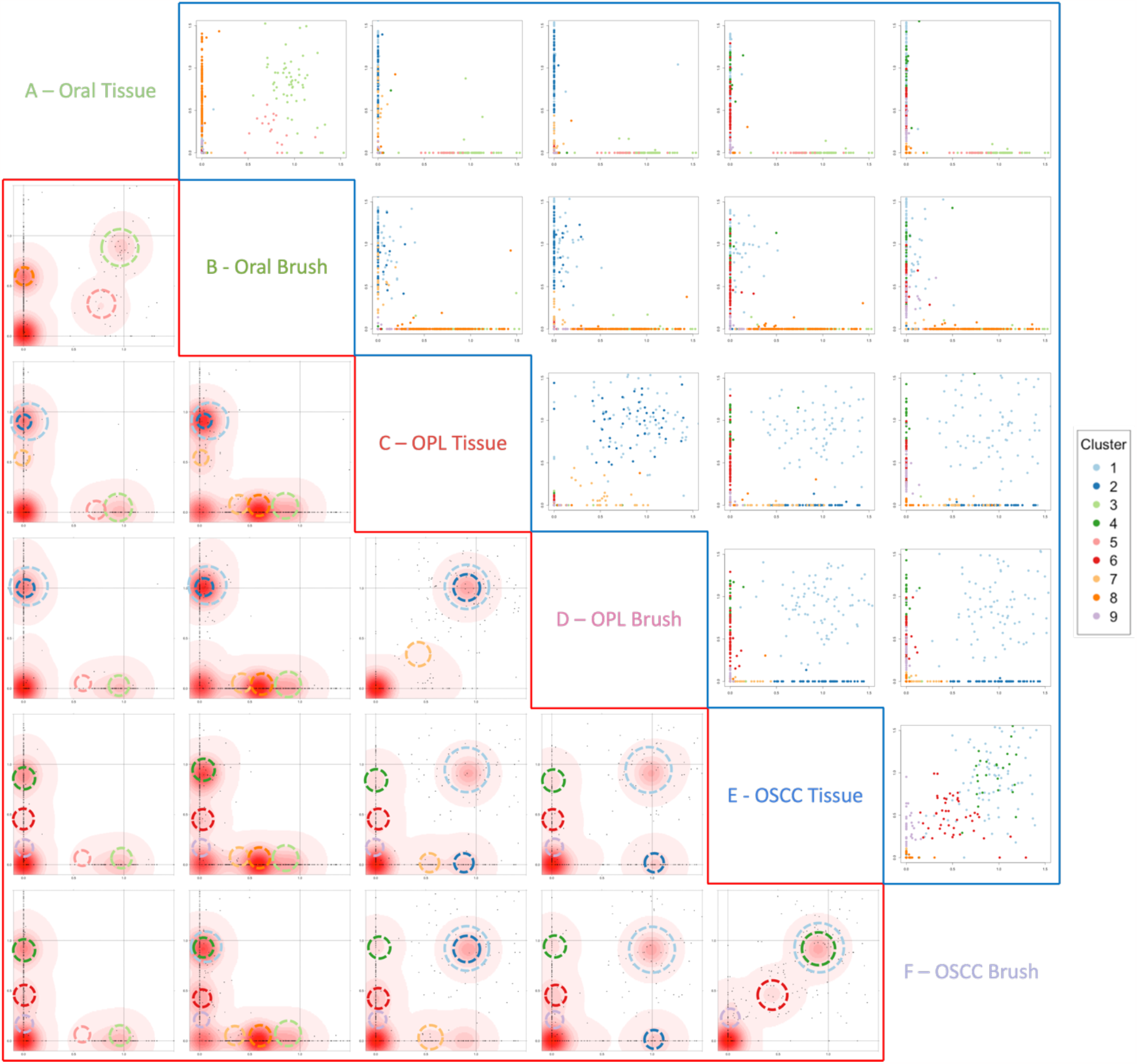
SNV Clustering using DPClust. Clusters are colored according to the legend provided. The bottom left red section shows the density of SNVs and the circles around the densities mark which color-coded cluster they belong to. The upper right blue section displays the distribution of the SNVs with each SNV colored according to the cluster it falls in.

**Figure 5:**
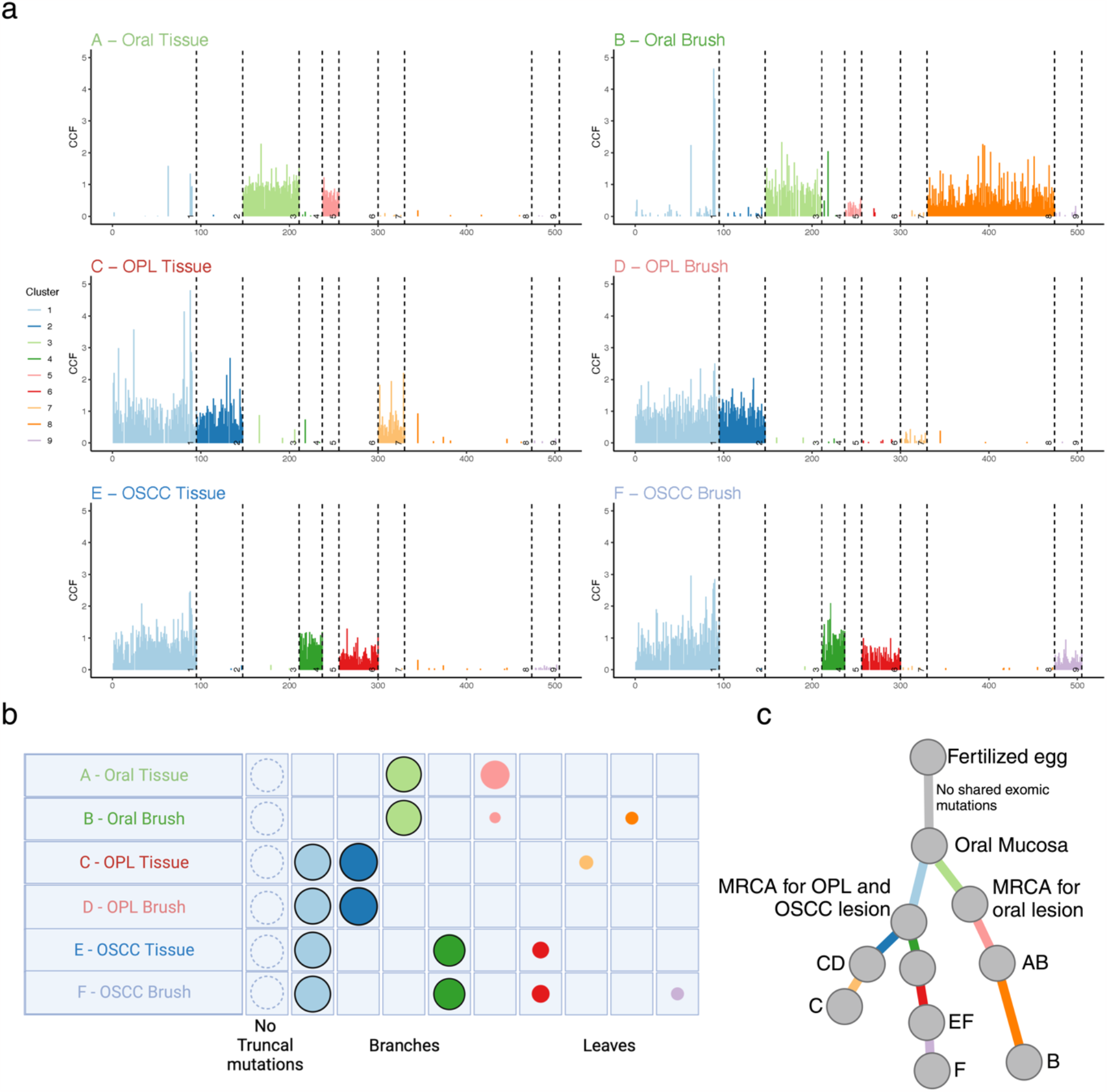
Reconstruction of subclonal architecture. (a) Line plot of CCF in each mutation. The plot shows the number of mutations in each cluster, clusters present in each sample, and the number of shared and unique clusters between samples. Each line represents a mutation, and the length of the line corresponds to the CCF value of that mutation. (b) Each clone and subclone detected is represented as a set of color-coded circles across all samples. Each row represents a sample, and the area of the circle is proportional to the CCF of the corresponding subclone. Clonal and near-clonal populations are shown with solid borders. The circle plots are divided into three types: trunk (CCF=1 in all samples, not seen in current samples), branches (present in more than one sample and either not found in all samples or subclonal in at least one), and leaf (specific to single samples). (c) Subclonal tree showing the relationship between subclones. The branch lengths of the phylogenetic tree are proportional to the number of mutations in each cluster and branches are annotated with samples in which they are present.

To assess the evolutionary history of the target lesions, a phylogenetic tree was constructed using the SNV-based clonal reconstruction (**Figure 5b and 5c**). This identified three distinct branches in the tree with an initial branching of OPL and OSCC together from the normal tissue through a shared common ancestor (branch AB and branch CDEF), consistent with the copy-number-based phylogenetic reconstruction (**Supplementary Figure 1**). Following this, there was a divergence between OSCC and OPL with unique subclones (branches CD and EF) (**Figure 5c**). It is important to note that the paired brush and tissue of each lesion consistently showed close phylogenetic relationships, denoting that brush biopsies can recapitulate the evolution of a tumor (**Supplementary Figure 2**).

To assess putative driver mutations in our clonal populations, the catalog of mutated cancer genes from the Interactive OncoGenomics (IntOGen)^14^ was used (**Figure 6**). There were distinct mutations in known driver genes amongst the identified clones. Most notably clonal cluster 1, which was shared between OPL and OSCC samples, had two *TP53* mutations (R267W and D352V). In addition to this, other potential driver mutations in known oncogenes and tumor suppressors were identified. These include *BCL2A* and *POLQ* in cluster 1, *PRDM14* and *STAG2* in cluster 2, *BCORL1* and *PCDH17* in cluster 4, and *LRP1B* in cluster 6. These mutations are involved in several important pathways, including tumor suppression, apoptosis and DNA repair^15^. This shows that each subclone harbors additional mutations which may impart a selective advantage.

**Figure 6:**
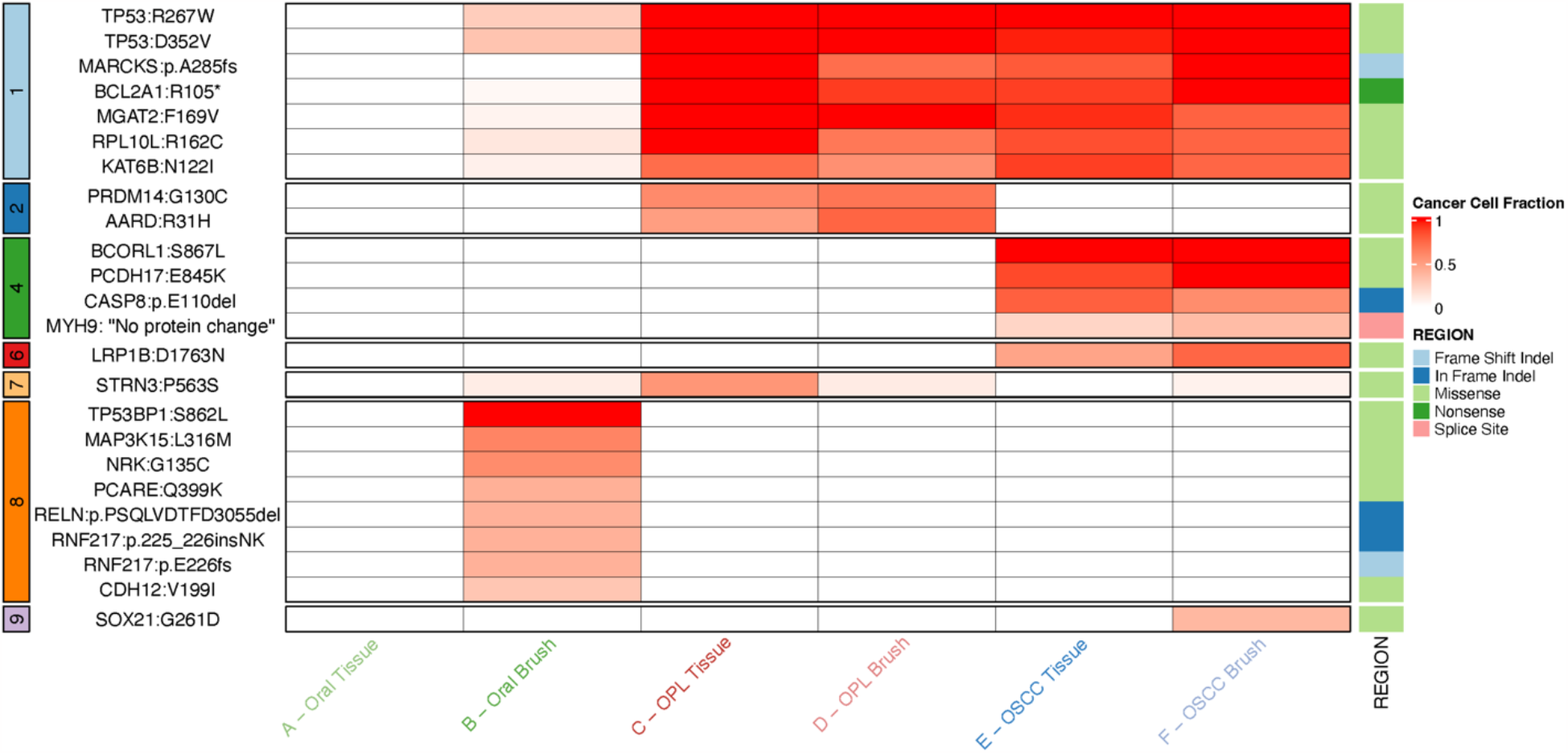
Oncoplot showing the presence of mutations in known driver genes in each sample, the cluster they fall in, and their CCF value. Genes and their protein change are denoted in each row. CCF values of the driver mutations are shown on a gradient scale from white to red for 0 to 1 CCF values. CCF of 0 (white) shows the absence of the mutation in the sample. The mutations are also denoted according to the cluster they fall in (left of the plot) and the type of mutation (right side of the plot).

This pilot proof-of-concept study successfully demonstrated that brush biopsies are effective in accurately replicating single nucleotide variants, copy number alterations, and phylogenetic relationships, in comparison to their paired tumor tissue. Moreover, our study confirmed the specificity of brush biopsies to their targeted tumor tissue. Indeed, tumor-specific alterations were not detected in adjacent normal tissue, while the number of mutations called was larger than in biopsies. The results highlight the potential of brush biopsy as a minimally invasive method of analyzing genomic alterations in OPLs, which could assist in identifying high-risk genomic alterations and subclones in the early stages of oral carcinogenesis, with the ultimate goal of enhancing molecular diagnosis and treatment.

## Methods

### Sample processing

We analyzed paired brush and tissue collected from a patient with both an OPL, diagnosed as moderate dysplasia, and oral squamous cell carcinoma, found on the contralateral sides of the tongue. In total, 6 samples were collected which were paired tissue and brush of the OPL, OSCC, and normal oral mucosa. Blood was also collected as the matched normal reference sample.

### Data analysis

#### Read mapping and quality control

Reads were mapped to the human reference genome (GRCh38) using the BWA-MEM algorithm (v.0.7.17)^16^ with default parameter as described in the DNA-Seq analysis Pipeline in NCI-GDC Documentation^17^. Quality control was performed using FastQC (v.0.11.5), SamTools (v.1.15)^18^, Picard (v.2.23.8)^19^, and Genome Analysis Toolkit (v.4.2.4.0)^20^. The files were recalibrated using BaseRecalibrator^11^ and ApplyBQSR^20^.

#### Somatic Mutation Calling

All samples were analyzed against the matched normal blood sample. Single Nucleotide Variants (SNVs) were called using Mutect2^20,21^ in tumor-normal mode. SNVs were filtered using the following cutoffs: (1) mapping quality score ≥50, (2) >2 tumor reads supporting variant, (3) tumor sequence depth at locus ≥10. Mutations were further visualized using the Integrative Genome Viewer^22^ for false positives. The filtered variants were annotated using Funcotator^11^.

#### Mutation Signature Analysis

We used SigFit^23^ to identify the linear combination of predefined Head and Neck Cancer signatures in COSMIC^24^ that most accurately reconstructed the mutational profile of the cohort. For the cohort signature analysis, signatures with contributions of less than 5% were assumed to be absent.

#### Copy number alteration calling

Segmental copy number information was derived for each sample using ASCAT (v3.1.2)^9^, as previously described, from which gains, losses, copy number-neutral events, and loss of heterozygosity (LOH) can accurately be determined. In addition, the ASCAT profiles reveal differences in aberrant tumor cell fraction, ploidy, gains, losses, LOH, and copy number-neutral events between the samples. The purity and ploidy estimates were cross-checked and if necessary refitted after DPClust clustering of mutation data and review of B allele frequency and LogR profiles. Minimum event distance (MED) based copy number phylogeny was constructed using MEDICC2^12^ to quantify the distance between copy number profiles.

### Subclonal reconstruction

To model the subclonal structure of the 6 oral samples sequenced, we employed DPClust^25,26^ in 6 dimensions. Indels were not considered in the clustering process, as their variant allele frequencies are over dispersed and biased towards the reference allele, which would add noise to the clustering. However, they were assigned to clusters post hoc based on their CCF values across the samples. Phylogenetic relationships between identified subclones were reconstructed following the pigeonhole principle^13^. 215 SNVs were excluded from the DPClust analysis since they had very low CCF in all of the samples (median CCF=0.0063), which is reminiscent of false positive artifacts.

## Acknowledgements

This work was funded by the Pre-Cancerous Atlas (PCA) at The University of Texas MD Anderson Cancer Center. T.M.B. is supported by a PhD fellowship from Boehringer Ingelheim Fonds. P.V.L. is a CPRIT Scholar in Cancer Research and acknowledges CPRIT grant support (RR210006).

## Author contributions

E.R.J. carried out the majority of the analyses with assistance from T.L. and supervision from X.Z. and P.V.L.; E.R.J., X.Z., and P.V.L. wrote the manuscript with comments and contributions from T.L., M.T., and J.R.W.; M.T. and T.M.B. provided bioinformatics assistance. A.G. provided the clinical samples and developed the clinical protocols. T.M.B., M.T., A.G. and J.R.W provided comments and edited the manuscript. All authors have read and approved the final manuscript.

## Ethics Declarations

The collection and analysis of tissues was approved at the UT MD Anderson Cancer Center under IRB protocols 2021-0129 and PA17-0050.

## Data availability

The datasets generated during and/or analyzed during the current study will be made available upon acceptance by journal.

## Figure legends

**Supplementary Figure 1** Copy number-based phylogenetic tree using MEDICC2. Pairwise distance was calculated based on the number of events. The oral epithelium samples (A and B) and the OSCC samples (E and F) were closer to each other.

**Supplementary Figure 2** Subclonal architecture reconstruction using only the same type of sample collection method. (a) Tissue biopsy and (b) brush biopsy. The branch lengths of the phylogenetic tree are proportional to the number of mutations in each cluster and branches are annotated with samples in which they are present.

