## Supplementary Figure 1 for "Reconstructing oral cavity tumor evolution through brush biopsy"

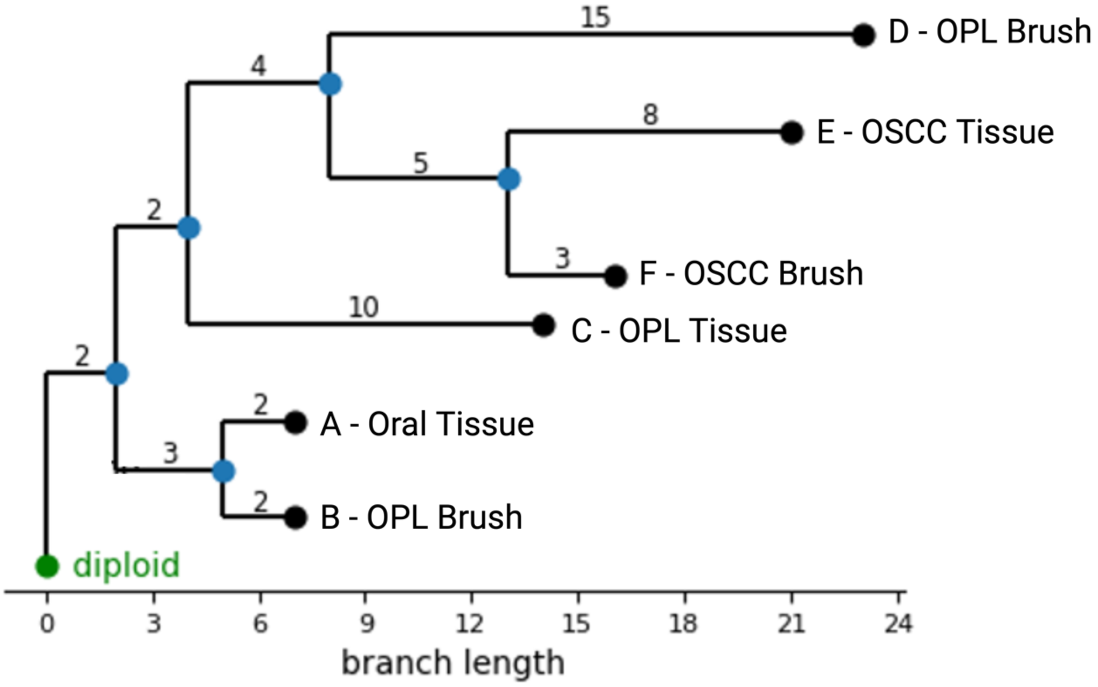


*Supplementary Figure 1: Copy number-based phylogenetic tree using MEDICC2 where pairwise distance was calculated based on the number of events. The oral epithelium samples (A and B) and the OSCC samples (E and F) were closer to each other compared to other lesions.*
