## Supplementary Figure 2 for "Reconstructing oral cavity tumor evolution through brush biopsy"

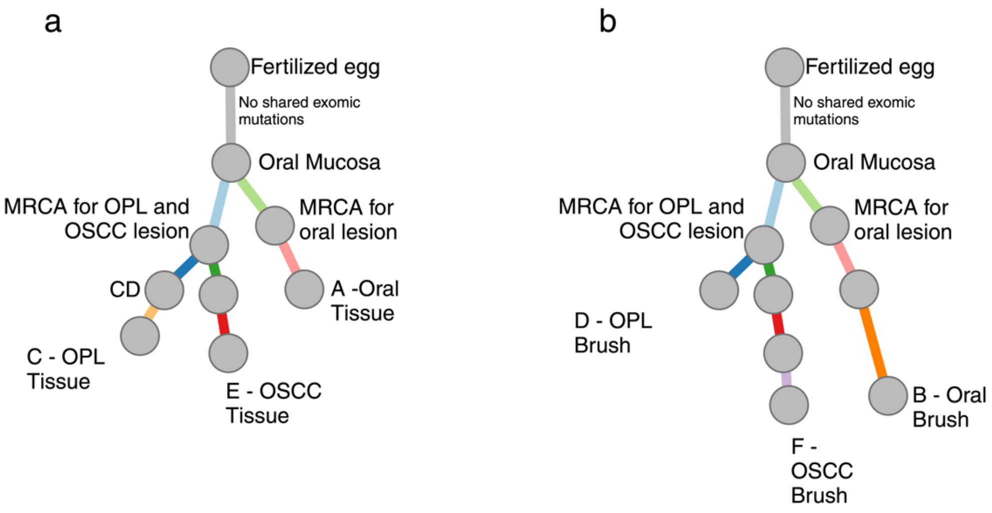


*Supplementary Figure 2: Subclonal architecture reconstruction using only the same type of sample collection method. (a) Tissue biopsy and (b) brush biopsy. The branch lengths of the phylogenetic tree are proportional to the number of mutations in each cluster and branches are annotated with samples in which they are present.*
